# The interaction of p38 with its upstream kinase MKK6

**DOI:** 10.1101/2020.11.26.395038

**Authors:** Ganesan Senthil Kumar, Rebecca Page, Wolfgang Peti

## Abstract

Mitogen-activated protein kinase (MAPK; p38, ERK and JNK) cascades are evolutionarily conserved signaling pathways that regulate the cellular response to a variety of extracellular stimuli, such as growth factors and interleukins. The MAPK p38 is activated by its specific upstream MAPK kinases, MKK6 and MKK3. However, a comprehensive molecular understanding of how these cognate upstream kinases bind and activate p38 is still missing. Here, we combine NMR spectroscopy and isothermal titration calorimetry to define the binding interface between full-length MKK6 and p38. We show that p38 engages MKK6 not only via its hydrophobic docking groove, but also helix αF, a secondary structural element that plays a key role in organizing the kinase core. We also show that, unlike MAPK phosphatases, the p38 conserved docking (CD) site is much less affected by MKK6 binding. Finally, we demonstrate that these interactions with p38 are conserved independent of the MKK6 activation state. Together, our results reveal differences between specificity markers of p38 regulation by upstream kinases, which do not effectively engage the CD site, and downstream phosphatases, which require the CD site for productive binding.

## 1. INTRODUCTION

The <u>m</u>itogen-<u>a</u>ctivated <u>p</u>rotein <u>k</u>inase p38α (MAPK14; α-isoform; 360 aa; 41.3 kDa; hereafter p38) has critical functions in cell differentiation, apoptosis and autophagy.^1^ Three MAPK Kinases (MAPKKs) phosphorylate p38 on a threonine and a tyrosine residue (p38α: T180 and Y182) in the p38 activation loop, which is necessary for p38 to be fully activated.^2^ The MAPKKs MKK3b and MKK6 are p38 specific, while MKK4 activates both p38 and JNK.^3,4,5^ Dephosphorylation by different downstream phosphatases, including the KIM-protein tyrosine phosphatases (KIM-PTPs), leads to p38 inactivation.^6,7^

Phosphatases, MAPKKs and substrates bind to p38 using short linear motifs (SLiMs) known as D-motifs or <u>K</u>inase <u>I</u>nteraction <u>M</u>otifs (KIM).^8,9^ Each KIM contains two to three basic residues, a short linker and a hydrophobic motif, Φ_A_-X-Φ_B_, where Φ_A_ and Φ_B_ are Ile, Leu, Met, or Val.^8^ The Φ_A_-X-Φ_B_ motif binds to a deep hydrophobic groove on the surface of p38 sandwiched between p38 helices αD-αE and the β7-β8 reverse turn.^10,11^ The basic residues of the KIM bind to complementary acidic residues in the so-called common docking (CD) site on p38.^9,12,13^ Structural studies of full-length proteins confirmed an extensive electrostatic interaction at the CD site between p38 and the KIMs of MK2, a substrate, and HePTP, a KIM-PTP.^14,15,16,17^

Previously, the structures of the complexes of p38 bound to the KIM peptides of MKK3b (MKK3bKIM; residues 16-32) and MKK6 (MKK6_KIM_; residues 4-17) were determined using X-ray crystallography (PDB IDs: 1LEZ, 2Y8O).^10,18^ While the MKK3/6 KIM sequences differ, with distinct flanking residues, the structures bound to p38 are essentially identical (RMSD = 0.14 Å). As expected, the KIM peptides bind the KIM binding pocket of p38. Unexpectedly, the p38 CD binding site was not occupied by the basic residues of MKK3bKIM and MKK6KIM, suggesting that either the electrostatic interaction between the CD site and the MKK peptide residues does not occur or that an electrostatic interaction occurs but is too weak and/or transient to be observed in the crystal structure. Furthermore, no molecular data has been reported for a full-length MKK:p38 complex.

Here we used biomolecular NMR spectroscopy and isothermal titration calorimetry (ITC) to address the following questions: (i) does the MKK6_KIM_ peptide interact with the p38 CD site in solution and (ii) is the interaction of p38 with full-length MKK6 different that of the MKK6_KIM_ peptide. We show that the MKK6_KIM_ peptide interacts extensively with the hydrophobic MAPK binding groove, but not with the CD site, consistent with the crystallographic results. Importantly, we discovered that full-length MKK6 interacts more extensively with p38 than the MKK6_KIM_ peptide, affecting also p38 helix αF, a secondary structural element that has previously been suggested to be important for the organization of activated p38.

## 2. RESULTS

MAPK p38 recognizes its upstream kinases, phosphatases and its substrates using the highly conserved D-/Kinase Interaction Motif (KIM). It has been previously shown that the upstream kinase MKK6 KIM motif peptide (MKK6_KIM;_ residues 4-17) binds p38 (K_D_ ~7 μM; SPR).^18^ We confirmed this result using ITC (K_D_ = 7 ± 2 μM) (**Supplementary Figure 1A; Supplementary Table 1**). Next, we employed NMR spectroscopy to define the molecular interactions of MKK6_KIM_ with p38. A direct comparison of the 2D [^1^H,^15^N] TROSY spectra of free and MKK6_KIM_-bound p38 reveals chemical shift perturbations (CSPs) of 28 peaks, 26 of which are in fast exchange and 2 peaks (L113 and N114) with line-widths broadened beyond detection (**Supplementary Figure 1B**). The majority of perturbations map to the p38 hydrophobic binding groove, affecting residues in helix αD, helix αE, the αD-αE loop, and the β7-β8 turn (**Supplementary Figure 1B, C**). No perturbations map to the p38 CD site residues. Together, these data confirm that the mode of interaction observed in the p38:MKK6_KIM_ crystal structure also occurs in solution.

Next, we investigated the interaction of full-length MKK6 with p38. When we previously compared the interaction of p38 with either KIM peptides or their corresponding full-length MAPK-phosphatases (HePTP, PTP-SL and STEP), interactions at the KIM site became stronger and additional CSPs were detected.^15,19^ The ITC data measured for the interaction of MKK6 and p38 is exothermic with a K_D_ of 2.0 ± 0.1 μM (**Figure 1A; Supplementary Table 1**), which is ~3.5 fold stronger than the interaction of MKK6_KIM_ with p38, suggestive of a larger binding interface between the two proteins. To identify the key residues involved in the interaction between p38 and MKK6, we formed the p38:MKK6 complex using size exclusion chromatography (SEC; **Supplementary Figure 1D**). A 2D [^1^H,^15^N] TROSY spectrum of the p38:MKK6 complex ((^2^H,^15^N)-labeled p38 and unlabeled MKK6) was then recorded and compared with the 2D [^1^H,^15^N] TROSY spectrum of free p38 (**Figure 1B**). The quality of the 2D [^1^H,^15^N] TROSY spectrum of the ~80 kDa p38:MKK6 complex is excellent and similar to the 2D [^1^H,^15^N] TROSY spectra recorded for other p38 complexes (p38:HePTP; p38:STEP).^15,19^ Direct comparison of the 2D [^1^H,^15^N] TROSY spectra with and without MKK6 reveals CSPs of 41 peaks, 30 in fast exchange and 11 peaks with linewidths broadened beyond detection (**Figure 1C**). Most of the perturbed residues belong to the hydrophobic docking groove of p38 (**Figure 1B, C, D**), as observed for MKK6_KIM_-bound p38. Residues that flank the p38 CD site also show small CSPs in the interaction with full-length MKK6. Finally, full-length MKK6 caused 11 peaks to broaden beyond detection (L113, H126, V127, D161, C162, L164, W207, V209, C211, A214, and E215). The majority of the line-broadened p38 peaks map to the hydrophobic docking groove. Notably, multiple residues in helix αF, which are located immediately below the hydrophobic binding groove, are also line broadened (**Figure 1E**). p38 helix αF serves as the major organizing center for the kinase.^20,21^ Two residues (I212 and L216) in helix αF are part of the ‘catalytic spine’ (C-Spine; residues V38, A51, L156, A157, V158, I212 and L216), which connects the kinase core to the ATP binding pocket through the hinge residue M109 and the p38 regulatory spine (R-Spine; residues L75, L86, H148 and F169) (**Figure 1F**). Thus, while the lack of CSPs for p38 residues distal from the KIM binding site suggests there are no additional interactions between the catalytic domain of MKK6 and p38, MKK6 binding to the p38 KIM-binding site cause significant changes to the p38 kinase core by affecting helix αF (our attempts to mutate residues in the helix αF to determine their exact role in the binding with fulllength MKK6 were unsuccessful, as helix αF variants result in protein that is insolubly expressed, indicating that helix αF plays an important role in the proper folding of p38).

**FIGURE 1.**
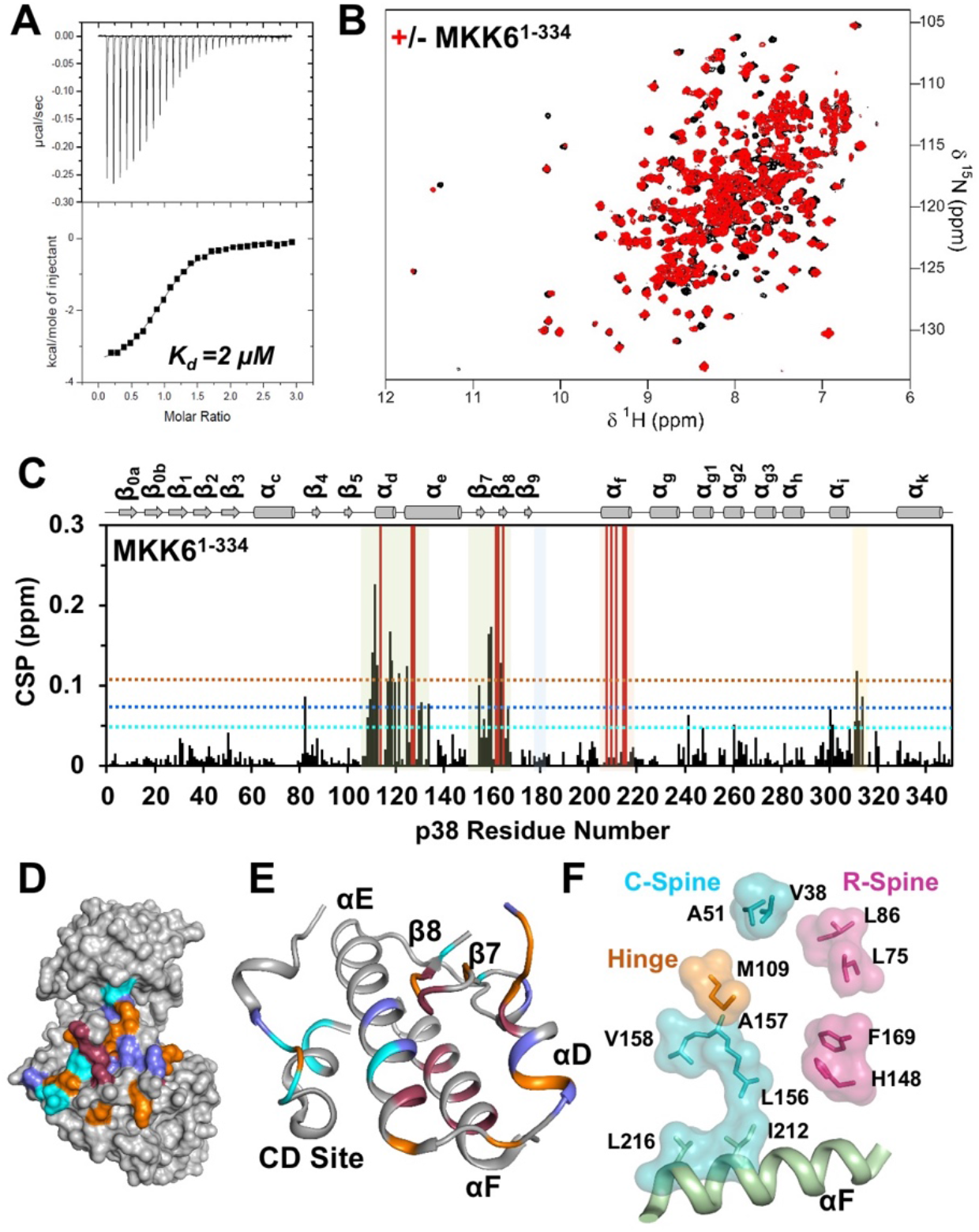
**(A)** Isothermal titration calorimetry of full-length MKK6 with p38. Data were recorded in triplicate. (**B**) Overlay of 2D [^1^H,^15^N] TROSY spectrum of (^2^H,^15^N)-p38:MKK6 (red) and (^2^H,^15^N)-p38 (black). (**C**) Histogram showing the ^1^H/^15^N chemical shift perturbations (CSPs) upon full-length MKK6 binding. Also highlighted are the key regions of p38, including the hydrophobic binding groove (green), the activation loop (blue), helix αF (orange) and the CD site (yellow). Peaks that are broadened beyond detection are indicated by red bars. The horizontal dotted lines correspond to 1σ (cyan), 2σ (blue) and 3σ (orange) chemical shift changes. (**D**) Surface representation of p38 residues that exhibit chemical shift perturbations (1σ-cyan; 2σ-blue; 3σ-orange; line broadened – raspberry) in the presence of full-length MKK6 *(bottom)* are mapped onto the p38 structure (PDB ID 5UOJ). (**E**) Cartoon representation showing the location of key regions involved in the interaction with MKK6 – hydrophobic binding groove (helix αD, αE, loop αD – αE, β-turns β7 and β8), CD site and helix αF. Note that the helix αF is sandwiched between the hydrophobic binding groove regions. Resides showing CSPs are color coded as in D. (**F**) Helix αF forms the central kinase core by anchoring the residues of catalytic spine. Residues L216 and I212 of helix αF are part of the catalytic spine (cyan), which connects the hinge residue M109 (orange) to the regulatory spine (pink).

In order to test the interaction of active MKK6 with p38, we repeated NMR experiments with the constitutively active MKK6 variant (MKK6_1-334_^S207ET211E^; henceforth, referred as mutMKK6; this MKK6 variant readily phosphorylates p38 *in vitro*^22^). The binding affinity of mutMKK6 with p38 is similar to that observed for MKK6 (3.5 ± 1.4 μM vs 2.0 ± 0.1 μM) (**Supplementary Figure 1A and Table 1**). Like MKK6, mutMKK6 also perturbed p38 residues clustered within the hydrophobic docking groove, helix αF and, to a lesser extent, the CD site (**Supplementary Figure 1C and D**). This shows that the p38 engages both inactive and active MKK6 in a similar manner.

## 3. DISCUSSION

Short linear motifs (SLiMs) are critical mediators of protein:protein interactions, especially for interactions involving kinases and phosphatases.^23^ The KIM SLiM is strictly required for the association of MAPKKs with MAPKs.^10,18^ However, the binding interface between p38 and a full-length MAPKK has never been molecularly characterized. Therefore, it was unknown if MAPKK binding affects residues distal from the p38 KIM-binding site. Here, we used NMR spectroscopy and ITC to define the binding interface of MKK6_KIM_, full-length MKK6 and activated MKK6 with p38. Consistent with published crystal structures, our solution-based data show that the MKK6_KIM_ does not interact with the p38 CD site. However, here, we also show that the interaction of p38 with full-length MKK6 is much more extensive than that with MKK6_KIM_, extending to helix αF and, to a lesser extent, the CD site.

Dynamic charged:charged interactions have recently been shown to be important for the interaction of intrinsically disordered regions (IDRs) of proteins with their cognate binding partners.^24,25,26^ The lack of a charged:charged interaction between the MKK6_KIM_ and the p38 CD site, and the limited CSPs observed with full-length MKK6 and the CD site, contrasts with the significant electrostatic interactions observed between the CD sites and the KIM-PTP family of phosphatases under identical experimental conditions.^15^ This finding is further supported by the p38:MKK3b_KIM_ crystal structure, where no interaction between the MKK3b_KIM_ basic residues and CD site was observed, and a hydrogen-deuterium exchange (HDX) study, which showed minimal interactions between the MKK3b_KIM_ peptide and the p38 CD site.^10,27^ Additionally, point mutations within the CD site of another MAPK ERK2 did not inhibit the binding, cytosolic retention and phosphorylation of ERK2 by its activating kinase, MEK1, while the same mutations completely disrupted the interaction of ERK2 with PTPRR (a KIM-PTP).^28^ Therefore, the interaction of the cognate upstream kinases with both p38 and ERK2 are similar. Critically, the distinct interactions of these MAPKs with their upstream kinases versus their downstream phosphatases now explains various gain-of-function mutations, which occur in the CD site of MAPKs from *Saccharomyces* and *Drosophila*.^29,30^ Thus, it is likely that mutations within the MAPK CD site allow the upstream kinase to bind normally and subsequently phosphorylate and activate the MAPK, while preventing the downstream phosphatases from binding, dephosphorylating and inactivating the MAPK.^31^

Full-length MKK6 binding to p38 affected several residues in helix αF, which serves as the major organizing center of the kinase and anchors the C-and R-spines, which are connected to all structural motifs that are important for catalysis, including the catalytic loop, the P+1 loop, and the phosphorylation loop.^20,32^ Three residues in particular, W207, C211 and E215, which were perturbed by MKK6 form a docking site for the kinase Ala-Pro-Glu (APE) motif.^20^ The highly conserved APE motif stabilizes the activation segment by docking to helix αF. Additionally, the APE motif stabilizes the P+1 loop providing a means by which helix αF, and these residues in particular, defines substrate binding at P+1 site. Helix αF was not affected by binding any of the full-length KIM-PTPs.^15,19^ Therefore, this finding may highlight a mechanism by which docking of the activating kinase, MKK6, to the p38 KIM-binding site prepares the kinase for phosphorylation by MKK6, in addition to the role of dynamics in the activation of p38.^33,34,22^ Collectively, our results reveal striking differences between specificity markers of p38 regulation by upstream kinases and downstream phosphatases, which may be a key factor in their differential regulation.

## 4. MATERIALS AND METHODS

### Peptide and protein preparation

The MKK6_KIM_ peptide (SQSKGKKRNPGLKIPKEAF; residues 2-20) was custom synthesized and HPLC purified (>95% purity; Biosynthesis, Inc.). The expression and purification of human p38 (residues 2-349, α-isoform) was carried out as previously described.^22^ For NMR experiments, expression of uniformly [^2^H,^15^N]-labeled p38 was achieved by growing cells in D_2_O based M9 minimal media containing 1g/L of ^15^NH4Cl and 4g/L of D-glucose as the sole nitrogen and carbon sources, respectively. Multiple rounds (0%, 30%, 50%, 70% and 100%) of D_2_O adaptation was necessary for high yield protein expression.^35^

MKK6 (residues 1-334) was subcloned into RP1B with an N-terminal His6-tag and a TEV protease recognition sequence.^36^ The expression vector was transformed into BL21-(DE3) RIL *E. coli* cells (Agilent). Cultures were grown in Luria broth at 37°C with shaking at 250 rpm until reaching an optical density (OD_600_) between 0.6-0.8. Expression was then induced by the addition of 1.0 mM isopropylthio-β-_D_-galactoside (IPTG; Goldbio), and the cultures were grown for an additional 18-20 hours at 18°C. The cells were harvested by centrifugation (6000 *xg,* 15 min, 4°C) and stored at −80°C until purification. His_6_-MKK_61-334_ was initially purified using immobilized metal affinity chromatography (IMAC). Following overnight cleavage with TEV protease in 50 mM Tris pH 8.0, 500 mM NaCl (4°C), MKK6_1-334_ was separated from the TEV and the His6-tag using a second IMAC step. The protein was further purified using size-exclusion chromatography (SEC; Superdex 200 26/60) pre-equilibrated in ITC buffer (10 mM Tris pH 7.5, 150 mM NaCl, 0.1 mM EDTA, 0.5 mM TCEP) or NMR buffer (10 mM HEPES pH 7.4 or 6.8, 150 mM NaCl, 5 mM dithiothreitol [DTT]). MKK6_KIM_ peptide was solubilized in the NMR buffer immediately prior to the experiment. Constitutively active mutMKK6 (MKK6_1-334_^S207ET211E^) was generated by site-directed mutagenesis and purified as described above.

### NMR spectroscopy

All NMR experiments were performed at 298 K on a Bruker Advance IIIHD 850 MHz (^1^H Larmor frequency) spectrometer equipped with a TCI HCN z-gradient cryoprobe. NMR samples were prepared in NMR buffer containing 10% D_2_O (vol/vol). NMR spectra were processed and analyzed using Topspin 3.0/3.1 (Bruker, Billerica, MA) and NMRFAM-Sparky.^37^ Solubilized MKK6_KIM_ peptide was added to (^2^H,^15^N)-p38in 1:1, 1:3, 1:5 and 1:8 excess peptide ratios (100 μM p38) and 2D [^1^H,^15^N] TROSY spectra recorded. Complex of ^2^H,^15^N-labeled p38 with unlabeled MKK6_1-334_ and mutMKK6_1-334_ was generated by mixing equimolar ratios of the proteins followed by purification using SEC (Superdex 200 26/60, pre-equilibrated in NMR Buffer). Backbone amide chemical shift deviations were calculated: Δδ= √(0.5 ((δ_HN,bound_ -δ_HN,free_)^2^ + 0.04 (δ_N,bound_ – δ_N,free_)^2^)).

### Isothermal Titration Calorimetry

ITC experiments were performed at 25°C using a VP-ITC micro-calorimeter (Microcal Inc.). Titrant (10 μL per injection) was injected into the sample cell over a period of 20 seconds with a 250 second interval between titrations to allow for complete equilibration and baseline recovery. 28 injections were delivered during each experiment, and the solution in the sample cell was stirred at 307 rpm to ensure rapid mixing. To determine the thermodynamic parameters *(ΔH, ΔS, ΔG)* and binding constant *(K)* of the p38:MKK6_1-334_ interaction, p38 was titrated into MKK6_1-334_, and the data was analyzed with a one-site binding model assuming a binding stoichiometry of 1:1 using Origin 7.0 software. A nonlinear least-squares algorithm and the titrant and sample cell concentrations were used to fit the heat flow per injection to an equilibrium binding equation, providing values of the stoichiometry (*n*), change in enthalpy (*ΔH*), and binding constant (*K*). All data were repeated in triplicate.

## ACKNOWLEDGEMENTS

The authors thank Drs. Dana Francis and Dorothy Koveal for help with early steps of the project. This research was supported by grant RSG-08-067-01-LIB from the American Cancer Society to R.P. and by R01GM100910 from the National Institute of Health to W.P. This research is based in part on data obtained at the Brown University Structural Biology Core Facility, which is supported by the Division of Biology and Medicine, Brown University. The authors declare no conflict of interest.

## AUTHOR CONTRIBUTIONS

**Ganesan Senthil Kumar** – Conceptualization, formal analysis, investigation, writing-original draft, writing-review and editing, supervision; **Rebecca Page** – Conceptualization, funding acquisition, writing-review and editing; **Wolfgang Peti** – Conceptualization, formal analysis, investigation, project administration, supervision, writing-review and editing

## REFERENCES

1. Cuadrado A, Nebreda AR (2010) Mechanisms and functions of p38 MAPK signalling. Biochem J 429:403–417.

2. Moriguchi T, Kuroyanagi N, Yamaguchi K, Gotoh Y, Irie K, Kano T, Shirakabe K, Muro Y, Shibuya H, Matsumoto K, et al. (1996) A novel kinase cascade mediated by mitogen-activated protein kinase kinase 6 and MKK3. J Biol Chem 271:13675–13679.

3. Han J, Lee JD, Jiang Y, Li Z, Feng L, Ulevitch RJ (1996) Characterization of the structure and function of a novel MAP kinase kinase (MKK6). J Biol Chem 271:2886–2891.

4. Lin A, Minden A, Martinetto H, Claret FX, Lange-Carter C, Mercurio F, Johnson GL, Karin M (1995) Identification of a dual specificity kinase that activates the Jun kinases and p38-Mpk2. Science 268:286–290.

5. Dérijard B, Raingeaud J, Barrett T, Wu IH, Han J, Ulevitch RJ, Davis RJ (1995) Independent human MAP-kinase signal transduction pathways defined by MEK and MKK isoforms. Science 267:682–685.

6. Muñoz JJ, Tárrega C, Blanco-Aparicio C, Pulido R (2003) Differential interaction of the tyrosine phosphatases PTP-SL, STEP and HePTP with the mitogen-activated protein kinases ERK1/2 and p38alpha is determined by a kinase specificity sequence and influenced by reducing agents. Biochem J 372:193–201.

7. Critton DA, Tortajada A, Stetson G, Peti W, Page R (2008) Structural basis of substrate recognition by hematopoietic tyrosine phosphatase. Biochemistry 47:13336–13345.

8. Bardwell AJ, Abdollahi M, Bardwell L (2003) Docking sites on mitogen-activated protein kinase (MAPK) kinases, MAPK phosphatases and the Elk-1 transcription factor compete for MAPK binding and are crucial for enzymic activity. Biochem J 370:1077–1085.

9. Tanoue T, Adachi M, Moriguchi T, Nishida E (2000) A conserved docking motif in MAP kinases common to substrates, activators and regulators. Nat Cell Biol 2:110–116.

10. Chang CI, Xu B, Akella R, Cobb MH, Goldsmith EJ (2002) Crystal structures of MAP kinase p38 complexed to the docking sites on its nuclear substrate MEF2A and activator MKK3b. Mol Cell 9:1241–1249.

11. Zhou T, Sun L, Humphreys J, Goldsmith EJ (2006) Docking interactions induce exposure of activation loop in the MAP kinase ERK2. Structure 14:1011–1019.

12. Tanoue T, Maeda R, Adachi M, Nishida E (2001) Identification of a docking groove on ERK and p38 MAP kinases that regulates the specificity of docking interactions. EMBO J 20:466–479.

13. Barsyte-Lovejoy D, Galanis A, Sharrocks AD (2002) Specificity determinants in MAPK signaling to transcription factors. J Biol Chem 277:9896–9903.

14. ter Haar E, Prabhakar P, Prabakhar P, Liu X, Lepre C (2007) Crystal structure of the p38 alpha-MAPKAP kinase 2 heterodimer. J Biol Chem 282:9733–9739.

15. Francis DM, Różycki B, Koveal D, Hummer G, Page R, Peti W (2011) Structural basis of p38α regulation by hematopoietic tyrosine phosphatase. Nat Chem Biol 7:916–924.

16. White A, Pargellis CA, Studts JM, Werneburg BG, Farmer BT (2007) Molecular basis of MAPK-activated protein kinase 2:p38 assembly. Proc Natl Acad Sci U S A 104:6353–6358.

17. Sok P, Gógl G, Kumar GS, Alexa A, Singh N, Kirsch K, Sebő A, Drahos L, Gáspári Z, Peti W, et al. (2020) MAP Kinase-Mediated Activation of RSK1 and MK2 Substrate Kinases. Structure 28:1101–1113.e5.

18. Garai Á, Zeke A, Gógl G, Törő I, Fördős F, Blankenburg H, Bárkai T, Varga J, Alexa A, Emig D, et al. (2012) Specificity of linear motifs that bind to a common mitogen-activated protein kinase docking groove. Sci Signal 5:ra74.

19. Francis DM, Kumar GS, Koveal D, Tortajada A, Page R, Peti W (2013) The differential regulation of p38α by the neuronal kinase interaction motif protein tyrosine phosphatases, a detailed molecular study. Structure 21:1612–1623.

20. Kornev AP, Taylor SS, Ten Eyck LF (2008) A helix scaffold for the assembly of active protein kinases. Proc Natl Acad Sci U S A 105:14377–14382.

21. Kim J, Ahuja LG, Chao F-A, Xia Y, McClendon CL, Kornev AP, Taylor SS, Veglia G (2017) A dynamic hydrophobic core orchestrates allostery in protein kinases. Sci Adv 3:e1600663.

22. Kumar GS, Clarkson MW, Kunze MBA, Granata D, Wand AJ, Lindorff-Larsen K, Page R, Peti W (2018) Dynamic activation and regulation of the mitogen-activated protein kinase p38. Proc Natl Acad Sci U S A 115:4655–4660.

23. Diella F, Haslam N, Chica C, Budd A, Michael S, Brown NP, Trave G, Gibson TJ (2008) Understanding eukaryotic linear motifs and their role in cell signaling and regulation. Front Biosci 13:6580–6603.

24. Borgia A, Borgia MB, Bugge K, Kissling VM, Heidarsson PO, Fernandes CB, Sottini A, Soranno A, Buholzer KJ, Nettels D, et al. (2018) Extreme disorder in an ultrahigh-affinity protein complex. Nature 555:61–66.

25. Wang X, Garvanska DH, Nasa I, Ueki Y, Zhang G, Kettenbach AN, Peti W, Nilsson J, Page R (2020) A dynamic charge-charge interaction modulates PP2A:B56 substrate recruitment. Elife 9.

26. Hendus-Altenburger R, Wang X, Sjøgaard-Frich LM, Pedraz-Cuesta E, Sheftic SR, Bendsøe AH, Page R, Kragelund BB, Pedersen SF, Peti W (2019) Molecular basis for the binding and selective dephosphorylation of Na+/H+ exchanger 1 by calcineurin. Nat Commun 10:3489.

27. Lee T, Hoofnagle AN, Kabuyama Y, Stroud J, Min X, Goldsmith EJ, Chen L, Resing KA, Ahn NG (2004) Docking motif interactions in MAP kinases revealed by hydrogen exchange mass spectrometry. Mol Cell 14:43–55.

28. Tárrega C, Ríos P, Cejudo-Marín R, Blanco-Aparicio C, van den Berk L, Schepens J, Hendriks W, Tabernero L, Pulido R (2005) ERK2 shows a restrictive and locally selective mechanism of recognition by its tyrosine phosphatase inactivators not shared by its activator MEK1. J Biol Chem 280:37885–37894.

29. Brunner D, Oellers N, Szabad J, Biggs WH, Zipursky SL, Hafen E (1994) A gain-of-function mutation in Drosophila MAP kinase activates multiple receptor tyrosine kinase signaling pathways. Cell 76:875–888.

30. Brill JA, Elion EA, Fink GR (1994) A role for autophosphorylation revealed by activated alleles of FUS3, the yeast MAP kinase homolog. Mol Biol Cell 5:297–312.

31. Tárrega C, Blanco-Aparicio C, Muñoz JJ, Pulido R (2002) Two clusters of residues at the docking groove of mitogen-activated protein kinases differentially mediate their functional interaction with the tyrosine phosphatases PTP-SL and STEP. J Biol Chem 277:2629–2636.

32. Kornev AP, Taylor SS (2010) Defining the conserved internal architecture of a protein kinase. Biochim Biophys Acta 1804:440–444.

33. Akella R, Min X, Wu Q, Gardner KH, Goldsmith EJ (2010) The third conformation of p38α MAP kinase observed in phosphorylated p38α and in solution. Structure 18:1571–1578.

34. Nielsen G, Schwalbe H (2011) NMR spectroscopic investigations of the activated p38α mitogen-activated protein kinase. Chembiochem 12:2599–2607.

35. Peti W, Page R (2016) NMR Spectroscopy to Study MAP Kinase Binding to MAP Kinase Phosphatases. Methods Mol Biol 1447:181–196.

36. Peti W, Page R (2007) Strategies to maximize heterologous protein expression in Escherichia coli with minimal cost. Protein Expr Purif 51:1–10.

37. Lee W, Tonelli M, Markley JL (2015) NMRFAM-SPARKY: enhanced software for biomolecular NMR spectroscopy. Bioinformatics 31:1325–1327.

