## Supplementary Information for "The interaction of p38 with its upstream kinase MKK6"

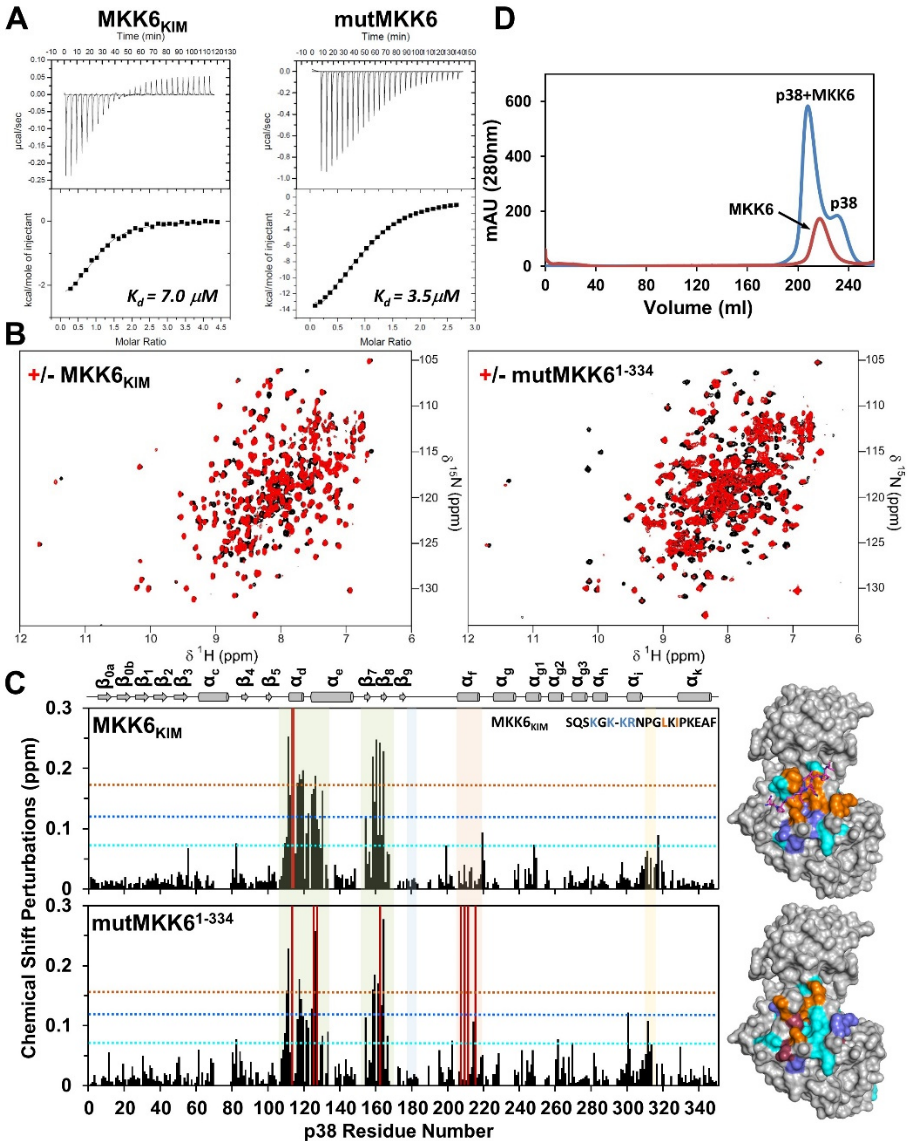


**SUPPLEMENTARY FIGURE 1. (A)** Isothermal titration calorimetry of MKK6$_{KIM}$ (*left*) and mutMKK6 (*right*) with p38. Data were recorded in duplicate. (**B**) Overlay of the 2D [$^{1}$H,$^{15}$N] TROSY spectrum of ($^{2}$H,$^{15}$N)-p38 in the presence (red) and absence (black) of MKK6$_{KIM}$ (*left*) and mutMKK6 (*right*). (**C**) Histogram showing the $^{1}$H/$^{15}$N chemical shift perturbations (CSPs) upon MKK6$_{KIM}$ and mutMKK6 binding. Also highlighted are the key regions of p38: hydrophobic binding groove (green), activation loop (blue), helix αF (orange), and CD site (yellow). Peaks that are broadened beyond detection are indicated by red bars. The horizontal dotted lines correspond to 1σ (cyan), 2σ (blue) and 3σ (orange) chemical shift changes. Also shown are the surface representation of p38 residues showing chemical shift perturbations (1σ-cyan; 2σ-blue; 3σ-orange; line broadened – raspberry) in the presence of MKK6$_{KIM}$ and mutMKK6 (*bottom*) are mapped onto p38 structure (PDB ID 5UOJ). (**D**) Size-exclusion chromatogram of p38:MKK6 complex performed using a Superdex S200 26/60 column. Note the change in the retention volume upon complex formation.

**Supplementary Table 1.** Thermodynamic and dissociation constants for p38:$MKK_{KIM}$ peptide and p38:MKK6 complexes derived from ITC experiments at 25°C.

| Peptide | $K_d$ (μM) | ΔG ($kcal \cdot mol^{-1}$) | ΔH ($kcal \cdot mol^{-1}$) | TΔS ($kcal \cdot mol^{-1}$) |
|---|---|---|---|---|
| **p38:$MKK6_{KIM}$** | 7 ± 2 | -7.0 ± 0.2 | -3.3 ± 1.1 | 3.8 ± 1.2 |
| **p38:MKK6** | 2 ± 0.1 | -7.8 ± 0.1 | -3.7 ± 0.1 | 4.1 ± 0.1 |
| **p38:mutMKK6** | 3.5 ± 1.5 | -7.7 ± 0.6 | -17 ± 0.8 | -9.3 ± 1.3 |

All experiments were conducted in triplicate.
